# Computational characterization of the interactions between novel Boronic Acid derivatives with urokinase-type plasminogen activator (uPA) and their binding energy

**DOI:** 10.1101/2023.06.05.543664

**Authors:** Syeda Mashaal Shah, Mehak Rafiq, Uzma Habib, Rehan Zafar Paracha, Maria Shabbir

**Affiliations:** School of Interdisciplinary Engineering and Sciences, National University of Sciences and Technology, Islamabad, Pakistan; Atta-ur-Rahman School of Applied Biosciences, National University of Sciences and Technology, Islamabad, Pakistan

**Author notes:** Corresponding author: Mehak Rafiq. These authors contributed equally to this work.

## Abstract

1.

Urokinase type plasminogen activator is expected to play a significant role in metastasis therefore various inhibitors are being prepared for this target protein. However, the binding site with residues that are involved in binding and inhibition is unidentified. Hence, comprehensive computational techniques are applied for finding the binding pocket, important amino acid residues and for the characterization of the binding energy of the best ligand among seven novel boronic acid derivative inhibitors within the binding pocket. Among seven test compounds, C_14_H_21_BN_2_O_2_S showed best results in structure based molecular docking through Molecular Operating Environment (MOE) and GOLD suit with −3.2481 kcal/mol binding affinity and 46.4523 GOLD Score. C_14_H_21_BN_2_O_2_S also showed high binding affinity within the binding pocket in DFT (Density Functional Theory) studies. DFT was carried out using hybrid functional B3LYP in combination with basis set LANL2DZ level of density functional theory on the extracted geometry of bound ligand C_14_H_21_BN_2_O_2_S to the binding pocket of uPA with a −2 charge on amino-acid residue ASP189. Computational analysis values on Geometric Optimization (opt), Single Point Energy (SPE) and Self-Consistent Reaction Field (SCRF) were 53.9, −66.3 and −49.0 respectively. Hence it is concluded that C_14_H_21_BN_2_O_2_S shows better binding with uPA binding pocket when there is a negative two charge on it ASP189 amino acid residue in the binding pocket. These seven ligands were also used for generating pharmacophore model through random selection with genetic algorithm by MOE having sensitivity of 79% towards the test set, specificity of 78% towards test set and 51% calculated Matthews coefficient correlation.

**Author Summary:** Boronic-acid based proteasome inhibitor like Bortezomib and Ixazomib are Food and Drug Administration (FDA) approved drugs, which are being used for fighting cancer. They can be considered as a template for understanding the pharmacokinetics and role of Boronic-acid ligands in the process. Boron-based warheads with stabilised functionality along with reduced toxicity are beneficent therapeutically. We have utilized computational quantum mechanical techniques in predicting binding free energies for ligands and proteins in a solvent environment. Instead of providing precise estimations, these techniques are more suitable for prediction purposes. The main challenge is developing inhibitors for uPA sub-sites that have high selectivity, potency, and improved pharmacokinetic properties. We have used Molecular docking and ligand-based techniques to analyze the binding interactions between seven ligands and uPA. Among these ligands, C_14_H_21_BN_2_O2_S_ is identified as the most appropriate inhibitor based on scores and its interactions with specific receptor amino acid residues. Computational quantum mechanical studies are conducted using electron density and hybrid functional B3LYP to determine the binding energy. A pharmacophore model is designed to identify crucial descriptors and search for compounds that can effectively inhibit uPA. The model’s accuracy is assessed through QSAR analysis, which reveals favorable hydrogen bond donor and acceptor groups as well as aromatic hydrophobic rings in proximity to the ligands. The designed model demonstrates good sensitivity, specificity, and calculated Matthews coefficient correlation.

## 3. Introduction

Cancer metastasis is the start of last stages of tumorigenesis; it is the dissemination of diseased cells from the site of origin through detachment, followed by the movement towards other sites for invasion by blood vessels or lymphatic vessels. It is responsible for 90% mobidity of cancer patients(1), and new prognostic markers are needed to correctly predict whether a patient will develop metastases later (2). The trypsin-like serine protease (TLSP) urokinase-type plasminogen activator (uPA) may play a role in the propagation of transformed cancerous cells that cause invasive spread, metastasis, and angiogenesis in aggressive cancers like prostate, triple-negative breast, pancreatic, gastric, and colorectal (3).

ELISA and other methods show that uPA plays a major role in metastases (4). Results show high levels of uPA in tumour cells with poor patient prognosis and are seen in different aggressive cancers, making it the highest-graded biomarker for cancer metastases with the highest level of evidence (according to US National Cancer Institute scale of evidence, i.e. 1) in clinical application by performing pooled analysis with a sample of 8377 primary breast cancer patients (5). This biomarker is the best predictor of progression-free and overall survival in many tumours (6). Novel uPA inhibitors were developed due to a deep commitment to tumour suppression. After much research, antibodies and peptidomimetics inhibited metastasis and tumour growth in mice (7)(8)(9). These successes inspired researchers to find less hazardous and more effective drug-like molecules. After its role in angiogenesis and metastasis was discovered, researchers began searching for drugs that could inhibit uPA and its endogenous receptor (uPAR). Peptide aldehydes were initially utilised to block cellular proteases; however, bioavailability, stability, and medication toxicity failed due to off-target toxicity (10). In 2010, Julian Adams produced peptide boronates, which were more potent against uPA (11). Boron’s vacant p-orbital accepts a lone pair of oxygen on the binding pocket’s serine residue to increase potency (12). These compounds are known for their strong selectivity toward serine proteases because they engage with residues in the S3 and S4 binding subsites of enzymes like elastase and chymotrypsin to produce maximum inhibitory activity. Thrombin prefers the catalytic region’s P1 basic residues. Thus, leucine boronic-acid inhibitors do not interrupt action (12). First described by Xue in 2017, flavonoid inhibitors inhibit uPA molecularly. The phenolic hydroxyl ring of flavonoids binds to the S1 pocket with an IC50 of 7μM (13). Pharmacists want to highlight effective boronic acid inhibitors that generate reversible covalent connections with target proteins(11). In 2003, bortezomib became the first selective proteasome inhibitor to enter phase II clinical trials for several cancers and haematological malignancies. Considering recent successes, boronic acid and its derivatives may help design new medications

## 4. Results and discussion

### A. Binding pocket prediction

In the UniProt data displayed for the protein uPA, there is no clearly defined binding site. Literature up to this point has focused on certain amino acid residues (ASP189, SER190) that exhibit enhanced potency and selectivity for the ligands, thereby reducing off-target toxicity (13)(14)(15). Two open-source docking tools DoGsite scorer and RaptorX were used to dock the ligands in different orientations with the protein uPA. DoGsite scorer predicted nine binding pockets, and the second binding pocket (P 1) with the highest drug score of 0.67 was selected. It contained amino acid residues Alanine183 (ALA183), ALA184, ALA221, Arginine217 (ARG217), Aspartate189 (ASP189), ASP194, ASP223A, Cysteine191 (CYS191), CYS220, Glutamine192 (GLN192), Glycine (GLY216), GLY219, GLY226, Histadine99 (HIS99), Isoleucine17 (ILE17), Leucine181 (LEU181), Lysine223 (LYS223), LYS224, Proline225, (PRO225), Serine146 (SER146), SER190, SER214, Threonine147 (THR147), THR229, Tryptophan21 (TRP215), Tyrosine (TYR171), TYR172, Valine213 (VAL213), VAL227 and was suggested to be used in further studies as it presented favourable interactions with the ligand. The predicted binding pockets are visualised using an open-source user-sponsored 3D molecular structure visualisation tool PyMOL which is issued by Schrodinger.

**Figure 1.**
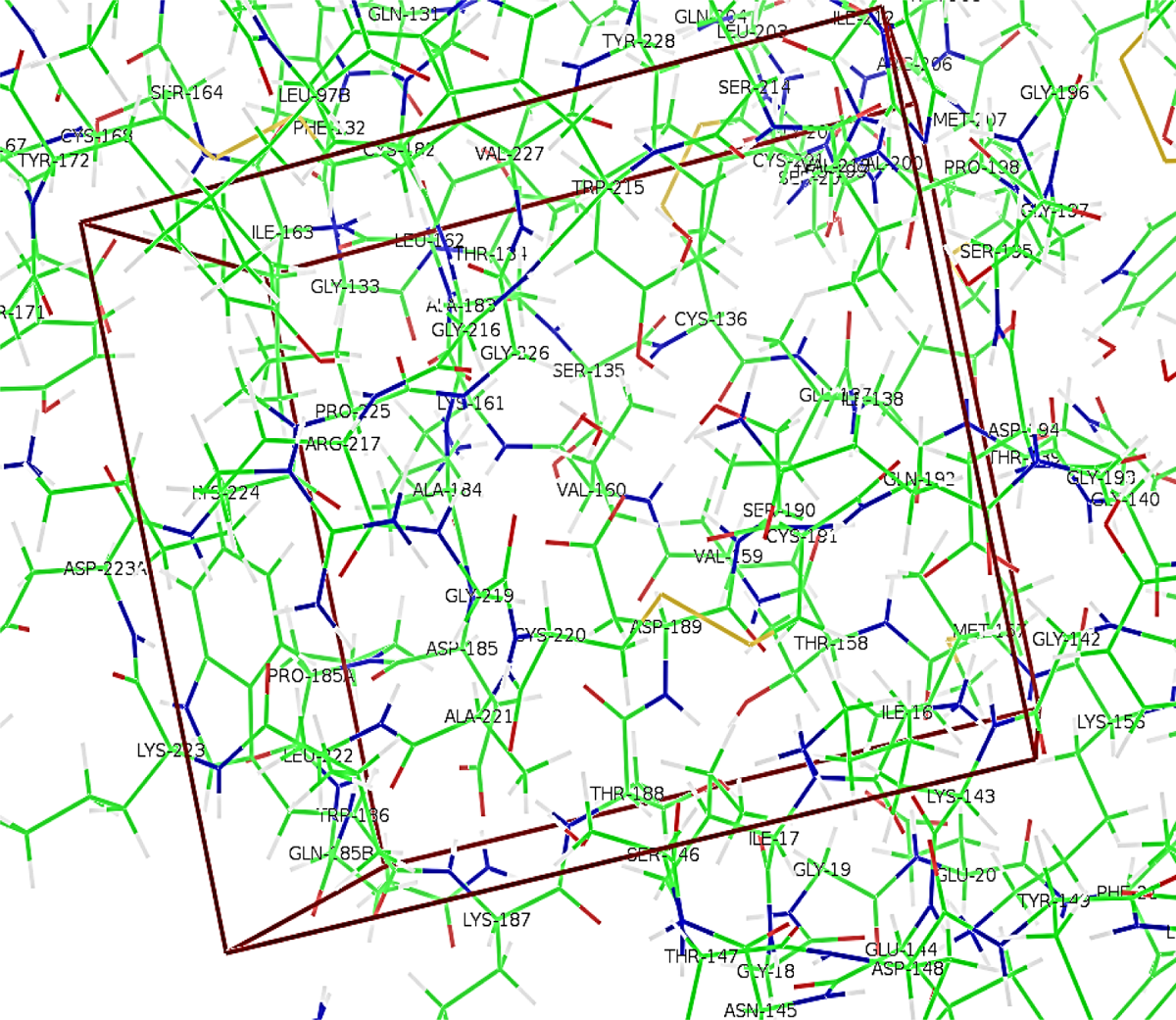
Binding pocket (P_1) predicted by DoGsite scorer. The amino acids contained in red outlined box represent the active binding site

For the validation of results obtained by the DoGSite scorer, RaptorX was used for the prediction of ligand binding pocket. RaptorX produced results showing four different binding sites for uPA for diverse ligands. Residues differ from each other for the generated binding pockets. The pockets are analysed based on their multiplicity, which is one of the confidence scores that indicate the goodness of the pocket based on the frequency with which the pocket appeared in the template structure. The first predicted pocket with the highest multiplicity 127 having the amino acid residues (H46, D192, S193, C194, Q195, G196, S198, V216, S217, W218, G219, G221, and C222) was most favourable for inhibition of uPA as suggested in the literature.

**Figure 2.**
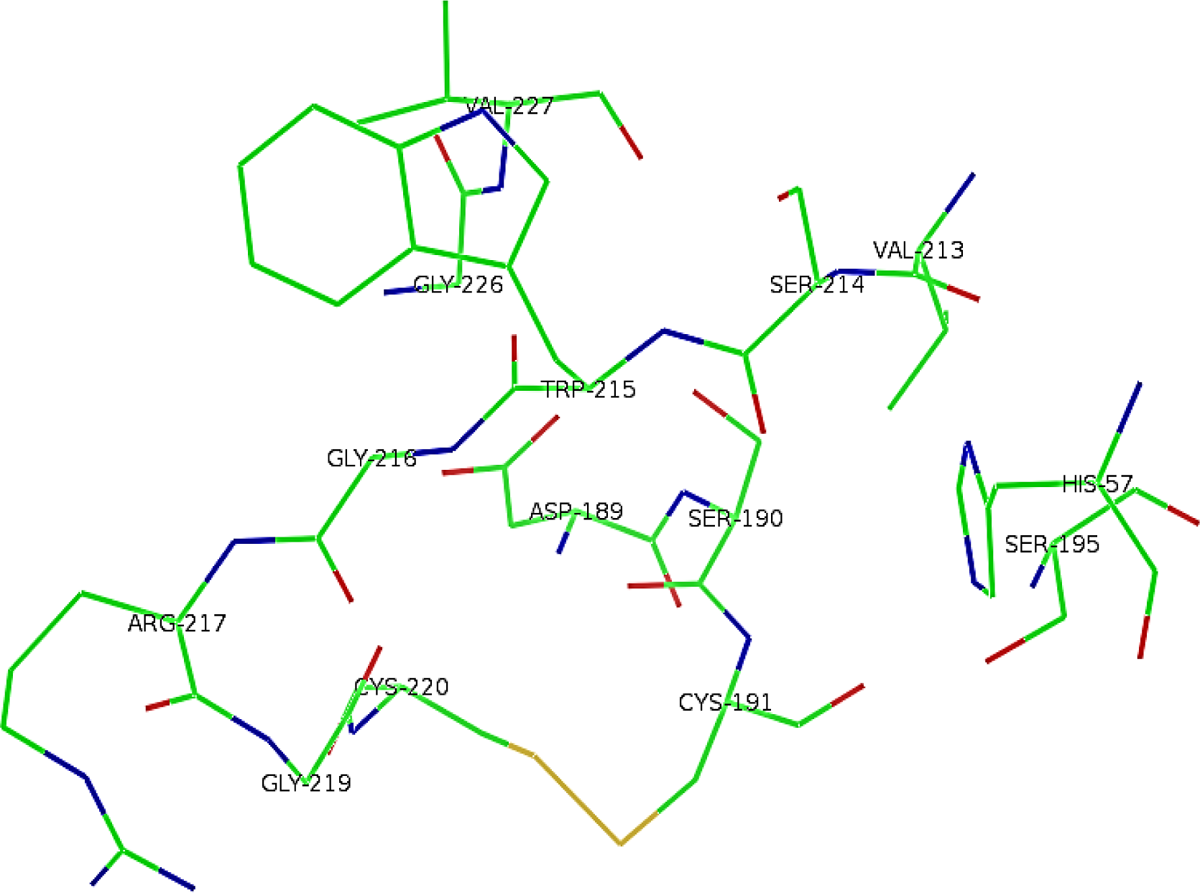
Selected binding pocket with multiplicity calculated as 127 generated with RaptorX. The animo acid residues depicted as sticks present in the pocket are labelled.

### B. Molecular docking studies

#### 1. Molecular docking studies using molecular operating environment (MOE)

While utilising the docking technique, the amino acid residues considered important in the binding pocket of uPA are as follows ASP189, SER190, and GLY219. For every ligand being studied, ten conformations were generated using the scoring function as LondonDG and the placement method as Alpha triangle. The interactions observed within the binding pocket were saved in the form of pictures with their binding affinity energy values given as scores in kcal/mol units. C_8_H_11_BN_2_O_2_S was docked in uPA with ten generated conformations. The third generated conformation showed the best score of −3.3591 kcal/mol, along with the best interactions within the binding pocket. One of the OH groups showed interaction with the residue ASP189, the Sulphur present in the ligand C_8_H_11_BN_2_O_2_S showed interaction with SER214 and NH_2_ group showed interaction with HIS57.

C_8_H_11_BN_2_O_2_S-a (right in Figure 3) was docked in uPA, and the first conformation showed the best score of −4.4899 kcal/mol, along with the best interactions within the binding pocket. The two OH groups showed interaction with the residues HIS57 and SER214, the NH_2_ group showed interaction with SER146 and GLN192, and the NH group of the ligand showed interaction with ARG217. For C₁₈H₃₁N₃O₂S (left in Figure 4), the first conformation showed the best score of − 5.3983 kcal/mol. Only the NH group showed interaction with ARG217 within the binding pocket. C_9_H_9_F_3_N_2_S (right in Figure 4) was docked, and the first conformation was selected as it gave the best score of −3.7709 kcal/mol. The Sulphur group showed interaction with the residue ASP189, and the NH_2_ group showed interaction with GLY219 and SER190.

**Figure 3.**
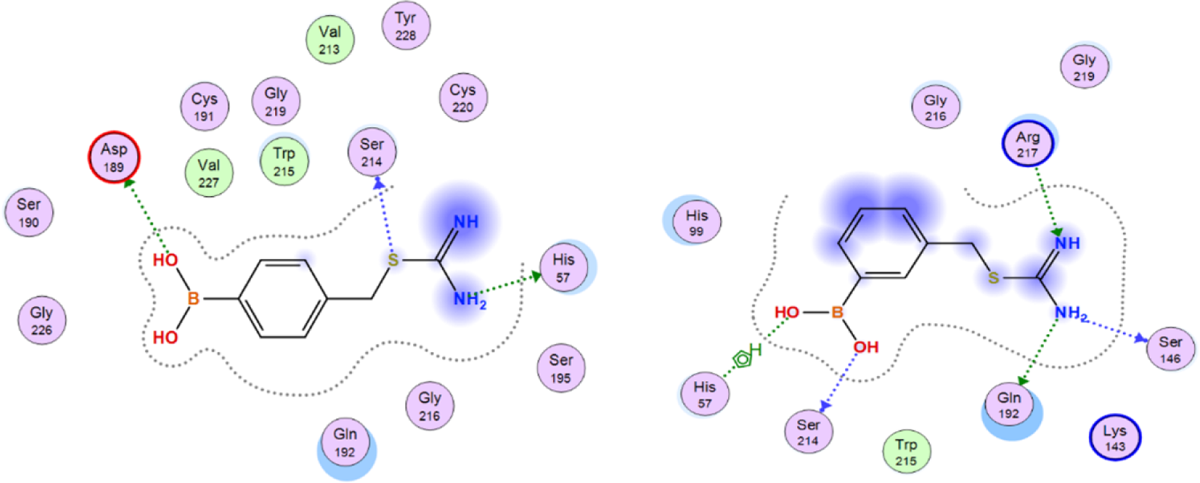
Visual representation of ligand *C_8_H_11_BN_2_O_2_S* (left) and *C_8_H_11_BN_2_O_2_S-a* (right) docked in uPA binding pocket.

**Figure 4.**
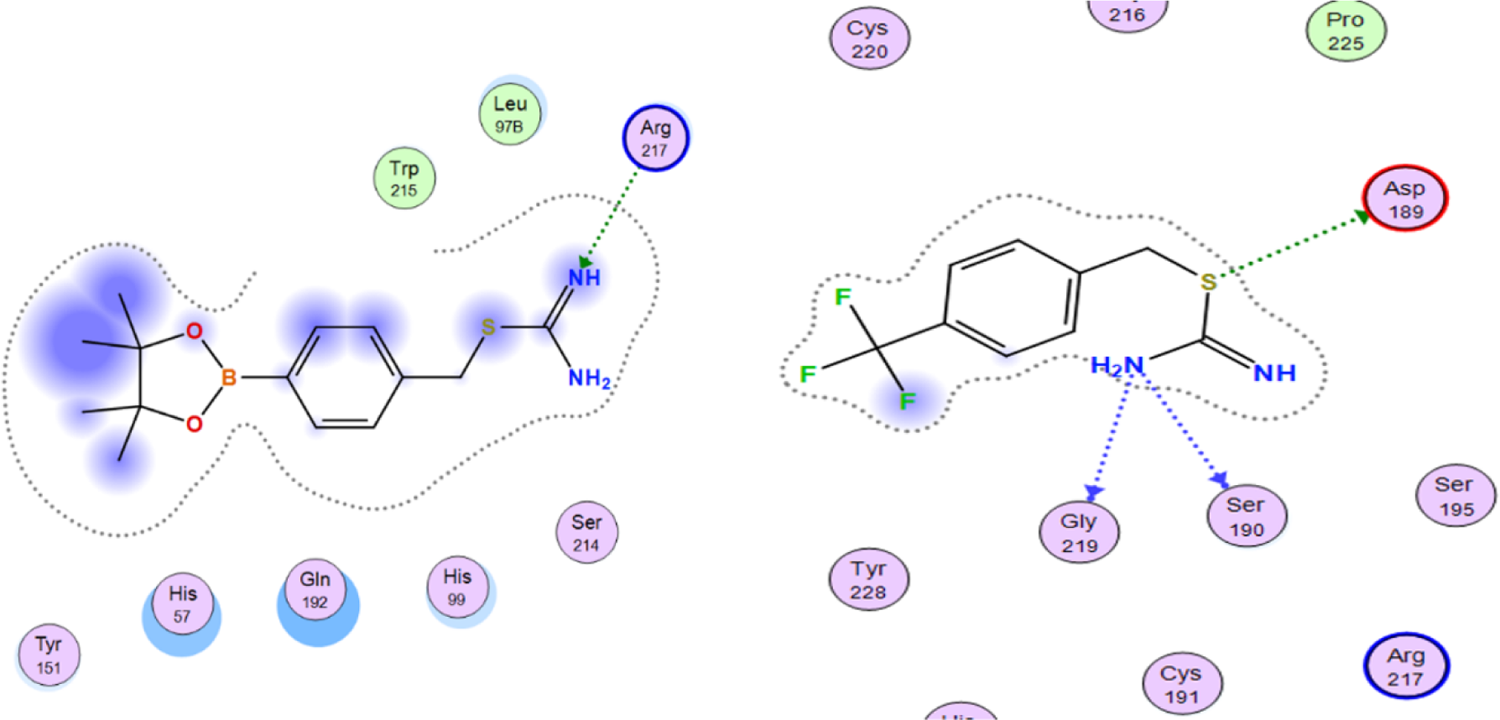
Visual representation of ligand *C₁₈H₃₁N₃O₂S* (left) and *C_9_H_9_F_3_N_2_S* (right) docked in uPA binding pocket.

C_14_H_21_BN_2_O_2_S (left in Figure 5) was docked in uPA with ten generated conformations. The first conformation showed the best score of −3.2481 kcal/mol, along with the best interactions within the binding pocket. The Sulphur present in the ligand C_14_H_21_BN_2_O_2_S shows interaction with GLY219, and the NH_2_ group shows interaction with ASP189 and SER190.

**Figure 5.**
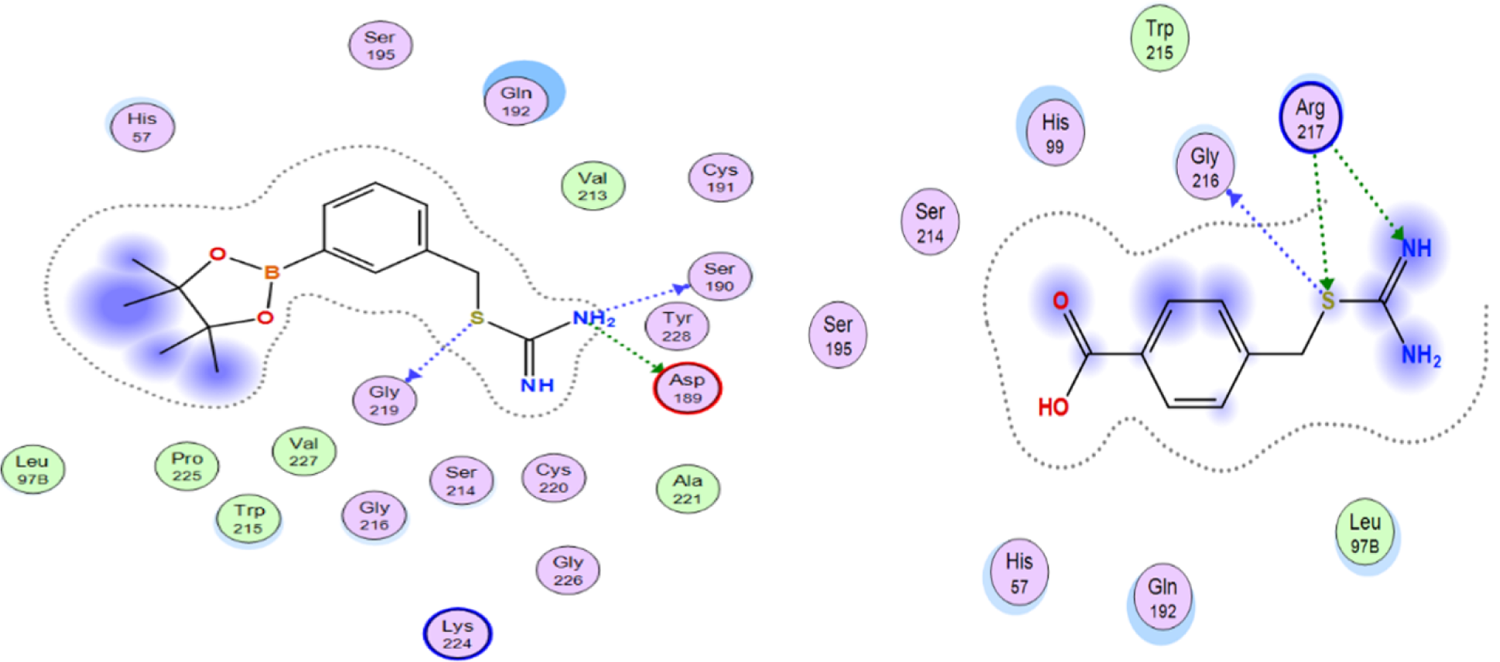
Visual representation of ligand *C_14_H_21_BN_2_O_2_S* (left) and *C₁₁H₁₈BNO₂S* (right) docked in uPA binding pocket.

C₁₁H₁₈BNO₂S (righ in Figure 5) was docked in uPA with ten generated conformations. The first conformation showed the best score of −4.8438 kcal/mol showing no favourable interactions within the binding pocket. The Sulphur present in the ligand C₁₁H₁₈BNO₂S shows interaction with GLY219 and ARG217, and the NH group also shows interaction with ARG217.

**Figure 6.**
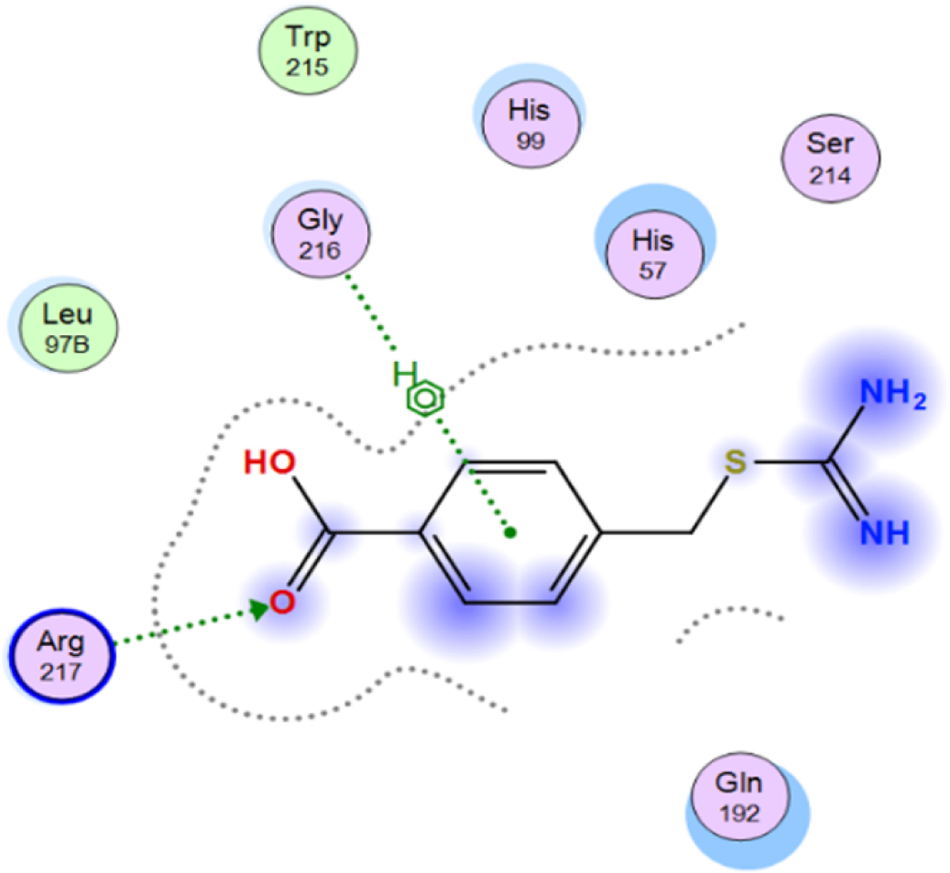
Visual representation of ligand *C_9_H_10_N_2_O_2_S* docked in uPA binding pocket.

C_9_H_10_N_2_O_2_S was docked in uPA, and the third conformation showed the best score of −4.6038 kcal/mol. The Oxygen functional group shows interactions within the binding pocket with ARG217, and the Benzene ring shows interactions with GLY216. Two compounds, C_8_H_11_BN_2_O_2_S and C_14_H_21_BN_2_O_2_S, were carefully chosen for further analysis for density functional theory studies as they showed favourable interactions within the binding pocket as a drug-like compound reveals activated only when it binds to its receptor-specific binding site. It was also revealed from previous wet lab studies by performing cell viability tests that four compounds, C₁₈H₃₁N₃O₂S, C_9_H_9_F_3_N_2_S, C₁₁H₁₈BNO₂S and C_9_H_10_N_2_O_2_S, caused cell death when Dictyostelium cells were subjected to acute and prolonged exposure to the test compounds. Although C_8_H_11_BN_2_O_2_S did not cause cell death, it did not show any favourable interactions within the binding pocket. Figure 12 provides the structures of the seven test compounds along with their scores, that is, the binding free energy (kcal/mol), electrostatic interaction energy (kcal/mol) and their van der wall interaction energy (kcal/mol) within the binding pocket of the protein of interest uPA.

#### 2. Molecular docking studies using GOLD Suit

Molecular docking was performed using a GOLD suit to verify the results obtained from MOE, and each pose was generated using the scoring function GOLD fitness score. Ten poses were generated for each test compound. The first conformation for C_8_H_11_BN_2_O_2_S (left in Figure 7) was suggested as the best fit solution with a score of 58.3489, along with favourable interactions within the binding pocket. The NH2 group showed interaction with ASP189 and Lus224, and the Sulphur atom present in the test compound C_8_H_11_BN_2_O_2_S showed interaction with TRP215. The second conformation for C_8_H_11_BN_2_O_2_S-a (right in Figure 7) was suggested as the best fit solution with a score of 65.1328, along with favourable interactions within the binding pocket. The NH_2_ group showed interaction with ASP189, ARG217, and LYS224, and the Sulphur atom present in the test compound C_8_H_11_BN_2_O_2_S-a showed interaction with TRP215.

**Figure 7.**
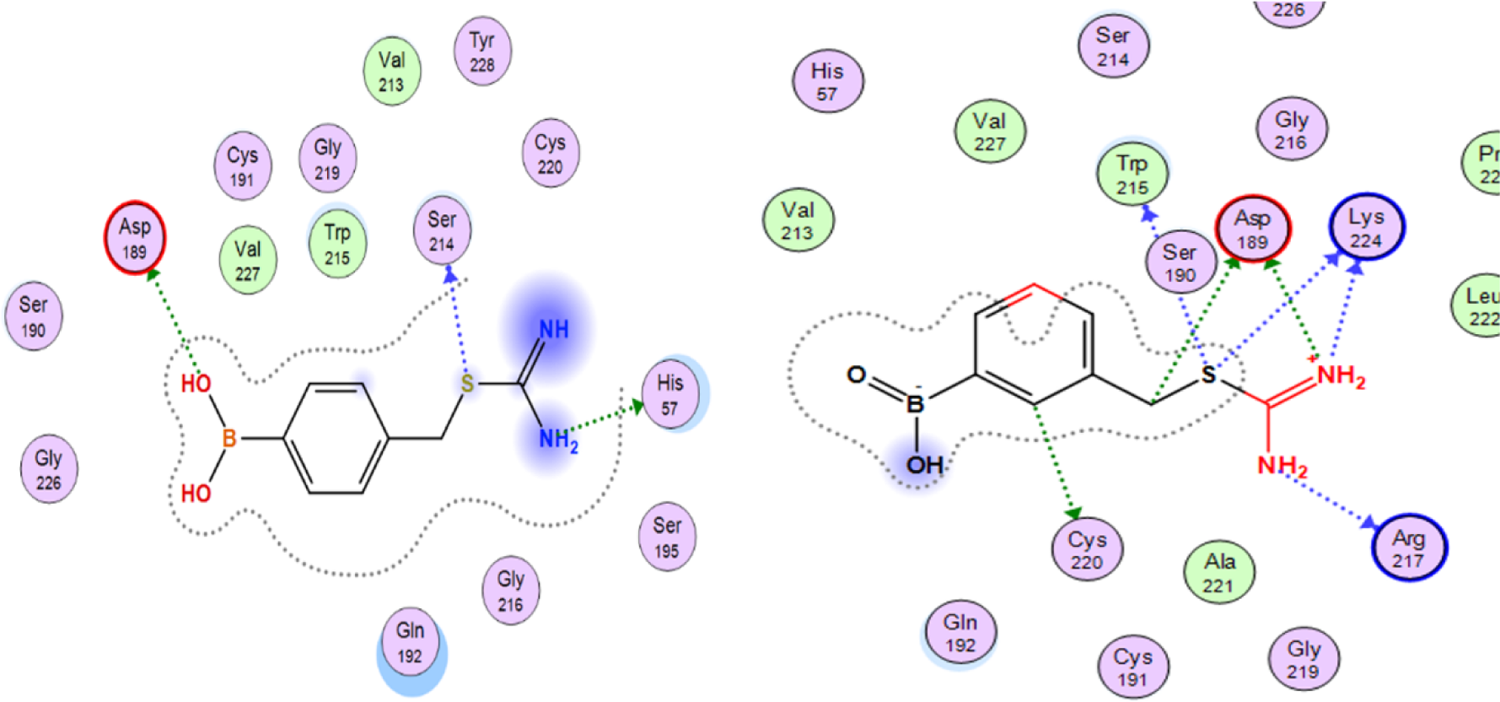
Visual representation of ligand *C_8_H_11_BN_2_O_2_S* (left) and *C_8_H_11_BN_2_O_2_S-a* (right) docked in uPA binding pocket.

The first conformation of C₁₈H₃₁N₃O₂S (left in Figure 8) was suggested as the best fit solution with a score of 47.9117, along with no favourable interactions within the binding pocket. The NH_2_ group showed interaction with TRP215. C_9_H_9_F_3_N_2_S (right in Figure 8) was docked in uPA, and the first conformation showed the best fit solution with a score of 57.0528, along with the best interactions within the binding pocket. The Sulphur present in the ligand C_9_H_9_F_3_N_2_S shows interaction with ASP189 and TRP215.

**Figure 8.**
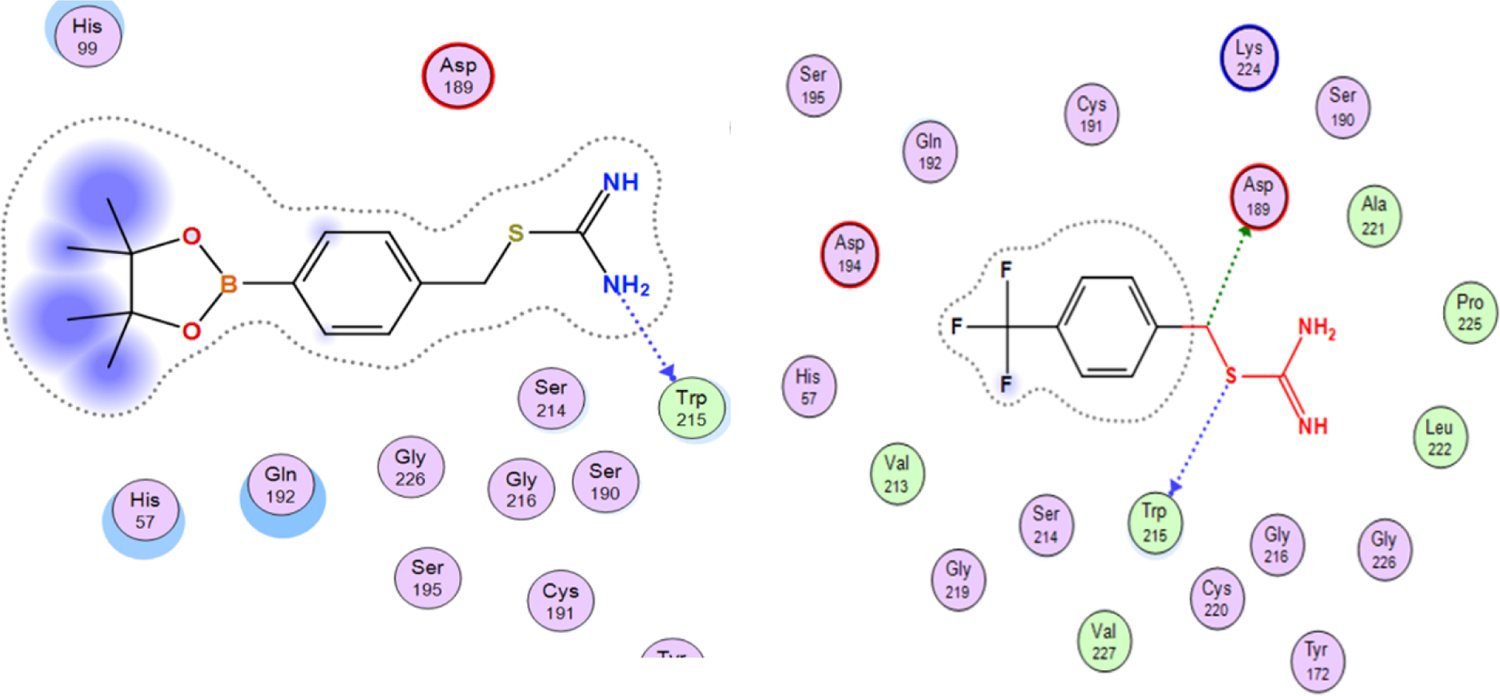
Visual representation of ligand *C₁₈H₃₁N₃O₂S* (left) and *C_9_H_9_F_3_N_2_S* (right) docked in uPA binding pocket.

C_14_H_21_BN_2_O_2_S (left in Figure 9) was docked in uPA with ten generated conformations, and the fourth conformation showed the best fit solution with a score of 46.4523, along with the best interactions. The Sulphur present in the ligand C_14_H_21_BN_2_O_2_S shows interaction with GLY219, the NH group shows interactions with the GLY216 of the binding pocket, and the NH_2_ group shows interactions with ASP189 of the binding pocket.

**Figure 9.**
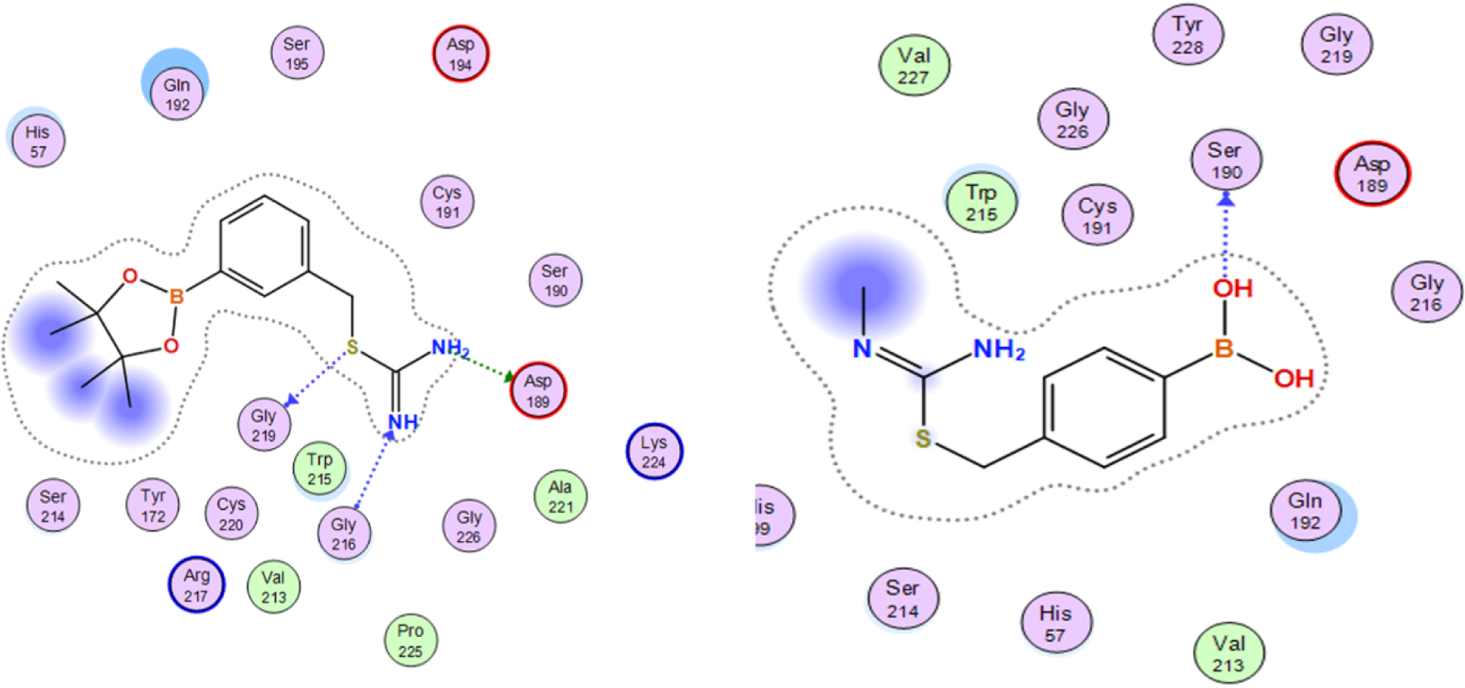
Visual representation of ligand *C_14_H_21_BN_2_O_2_S* (left) and *C₁₁H₁₈BNO₂S* (right) docked in uPA binding pocket.

**Figure 10.**
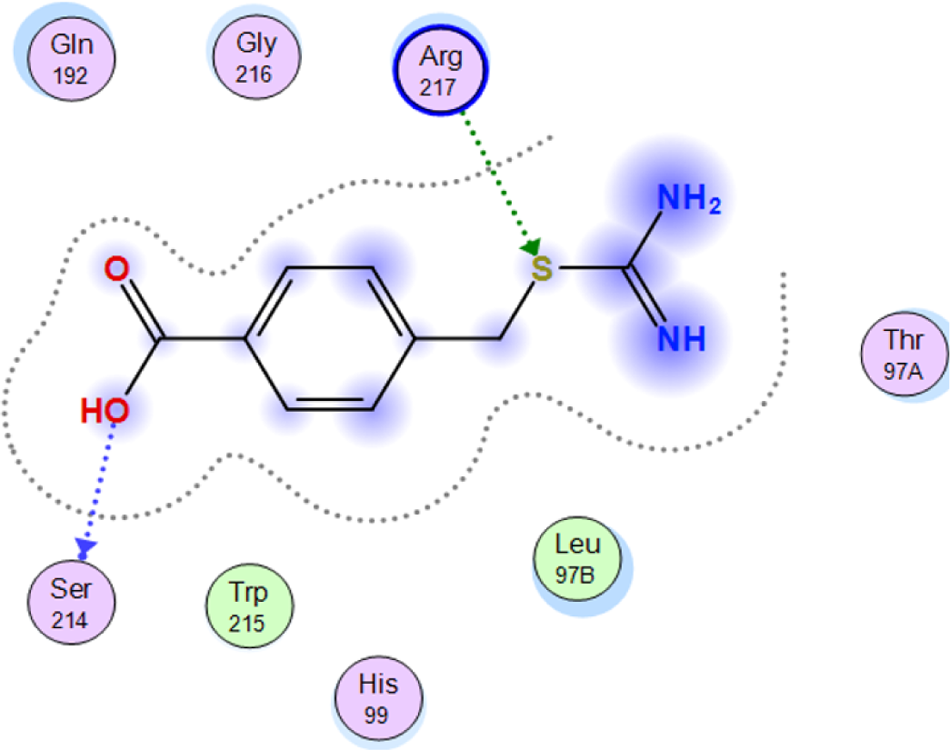
Visual representation of ligand *C_9_H_10_N_2_O_2_S* docked in uPA binding pocket.

For C₁₁H₁₈BNO₂S (right in Figure 9), the first conformation showed the best fit solution with a score of 53.7381. However, the test compound does not show any favourable interactions within the binding pocket. The OH present in the ligand C₁₁H₁₈BNO₂S shows interaction with SER190.

C_9_H_10_N_2_O_2_S was docked in uPA with ten generated conformations. The fifth conformation showed the best fit solution with a score of 21.2873. However, the test compound does not show any favourable interactions within the binding pocket. The OH present in the ligand C_9_H_10_N_2_O_2_S shows interaction with SER214, and Sulphur showed interactions with ArRG217.

### C. Quantum mechanical studies

#### 1. Model extraction

The ligand protein complex of uPA with its ligand C_14_H_21_BN_2_O_2_S obtained as a result through docking using GOLD software was further utilised for quantum mechanical studies. It is not computationally feasible to perform Density functional studies on the complete protein as it is very costly and would take a lot of time in calculating results. Therefore, the amino acid residues in close vicinity to the ligand (C_14_H_21_BN_2_O_2_S) were extracted for further evaluation using a free protein homology modelling server Swiss-Pdb (SPDB) Viewer. For quantum mechanical studies, the protein ligand complex needs to be truncated and reduced to the level of only ligand and the amino acid residues, ASP189, GLY216, GLY219, SER190, SER214, HIS57, TRP215 and LYS224 in the target protein located at the binding site and takes part in the binding interactions. The extracted model is given below in Figure 11

**Figure 11.**
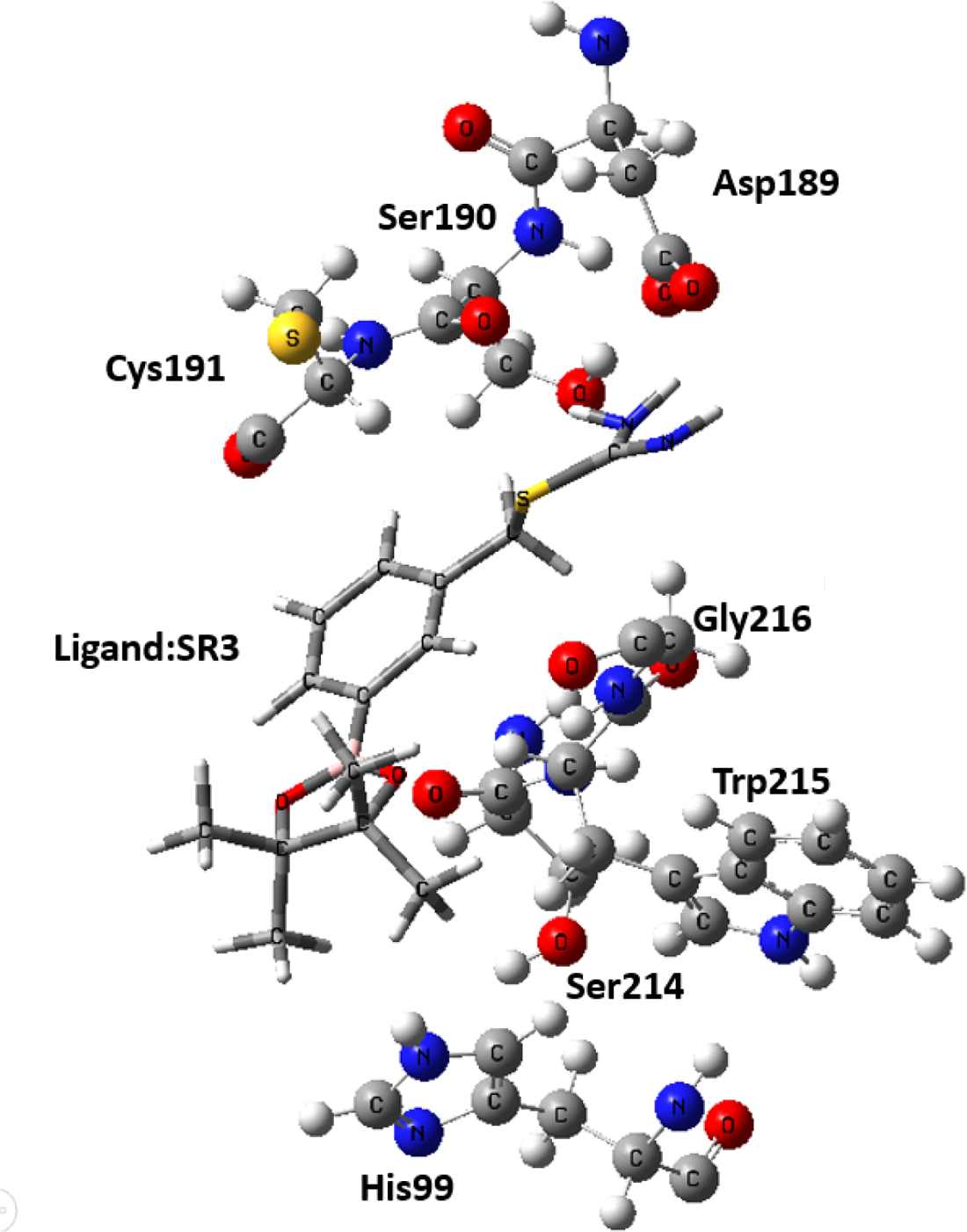
Selected residues of the binding pocket of uPA with its bound ligand extracted using SPDB viewer. The tube represents the ligand *C_14_H_21_BN_2_O_2_S* and the ball and stick model represent the amino acid residues of the binding pocket. For simplicity and clear visualise

#### 2. Ligand-protein complex geometry optimization

The extracted model of the binding pocket with bound ligand C_14_H_21_BN_2_O_2_S was passed through a series of clippings and modifications by trimming the amino acid residues ASP189, HIS99, SER214 and TRP215 at their alpha carbon (α-carbon) atom positions. To satisfy the valency where the cuttings were made, hydrogen atoms were added to the carbons of HIS99, SER214 and TRP215. Figure 12 given below depicts the selected binding pocket in complex with ligand C_14_H_21_BN_2_O_2_S with the needed clippings and modifications.

**Figure 12.**
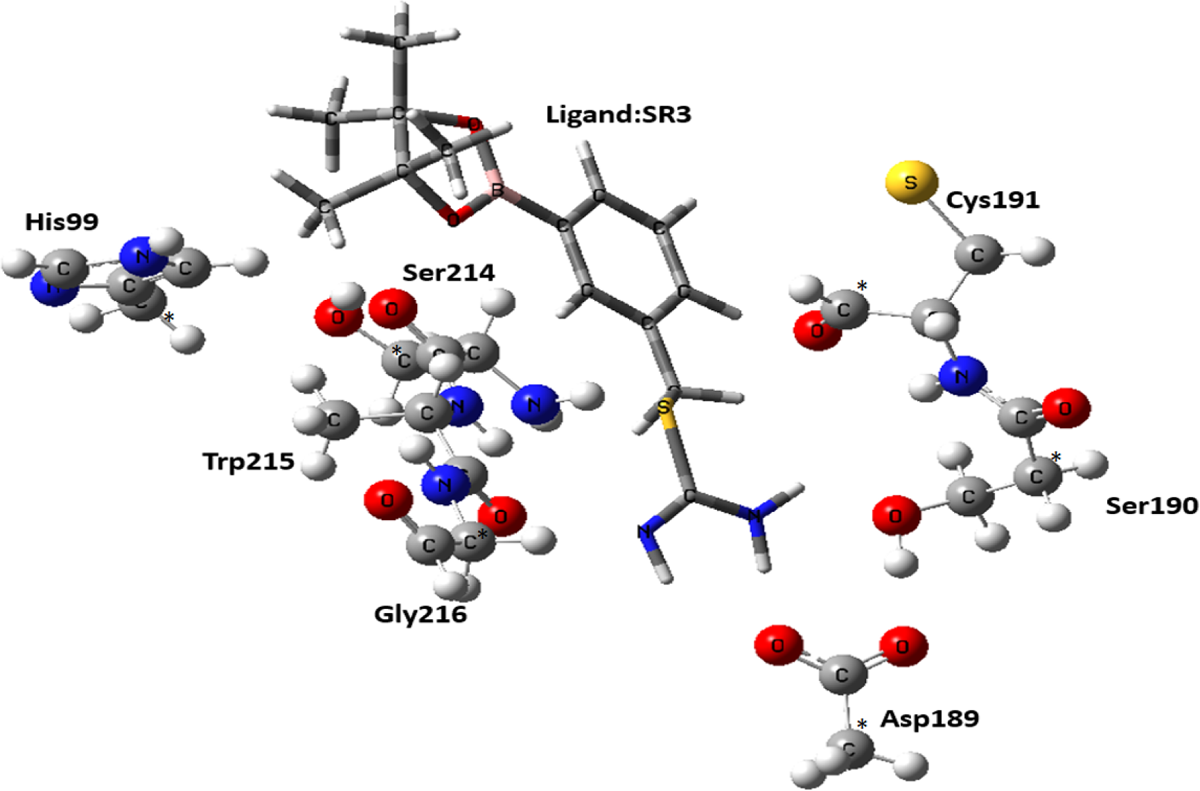
Proposed optimized model complex of uPA binding pocket with ligand *C_14_H_21_BN_2_O_2_S*. The sticks represent the ligand *C_14_H_21_BN_2_O_2_S* while the ball and lines represent the amino acid residues of the binding site. Asterics represent the fixed carbons where the cuttings are made.

The modified complex geometry of the product designed was subjected to optimization. The first step was hydrogen optimization where all the atoms except hydrogen were fixed. Once the hydrogen were optimized, this optimized geometry is then subjected to the next step that is geometry optimization of the model complex. During geometry optimization some of the atoms at the alpha carbons of amino acid residues ASP189, HIS99, SER214 and TRP215 are fixed, throughout the quantum mechanical studies being carried out, at locations where the trimmings were made so that they would stay on their positions of the X-ray crystal structure and retain the effect of normal protein.

For geometry optimization hybrid density functional method B3LYP was used in combination with a basis set LANL2DZ. The optimized geometry was then further utilised in the calculation of the single point energies in both gas and solvent phase using B3LYP/LANL2DZ level of DFT.

#### 3. Binding pocket geometry optimization

To understand the protein binding interaction energy, the optimized geometry of protein-ligand complex were used. To obtain minimized energy for protein binding pocket without ligand, ligand structure was removed and then protein binding pocket geometry model was optimized using same level of DFT, B3LYP/LANL2DZ. Figure 13 given below depicts the selected binding pocket without the ligand.

**Figure 13.**
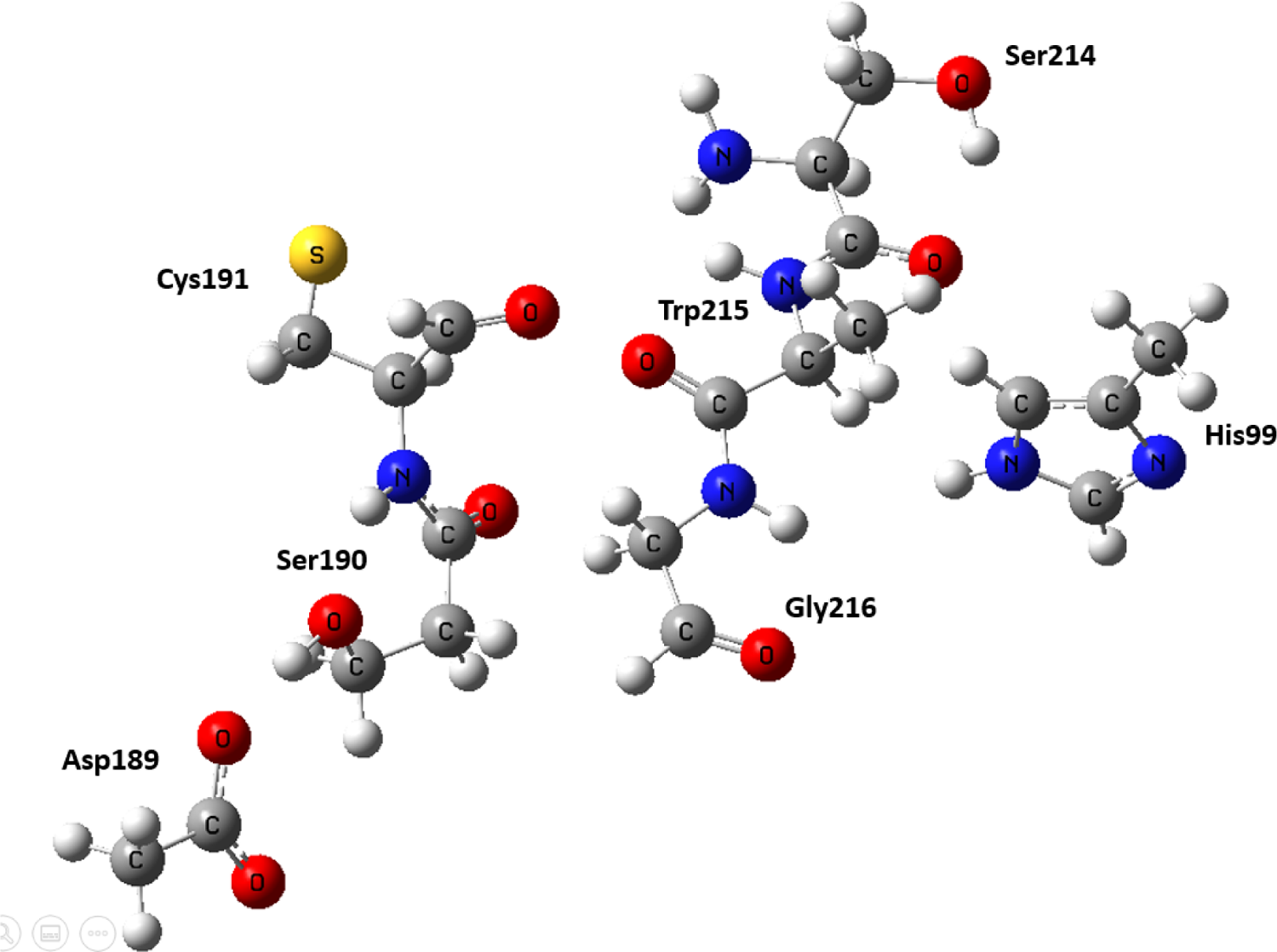
The optimized binding pocket of uPA

#### 4. Ligand Geometry optimization

For ligand model geometry optimization, ligand C_14_H_21_BN_2_O_2_S was extracted from X-ray crystal structure, PDB ID: 1W10 (16). Figure 14 shows the image of optimized ligand C_14_H_21_BN_2_O_2_S which was optimized using B3LYP/LANL2DZ level of DFT studies.

**Figure 14.**
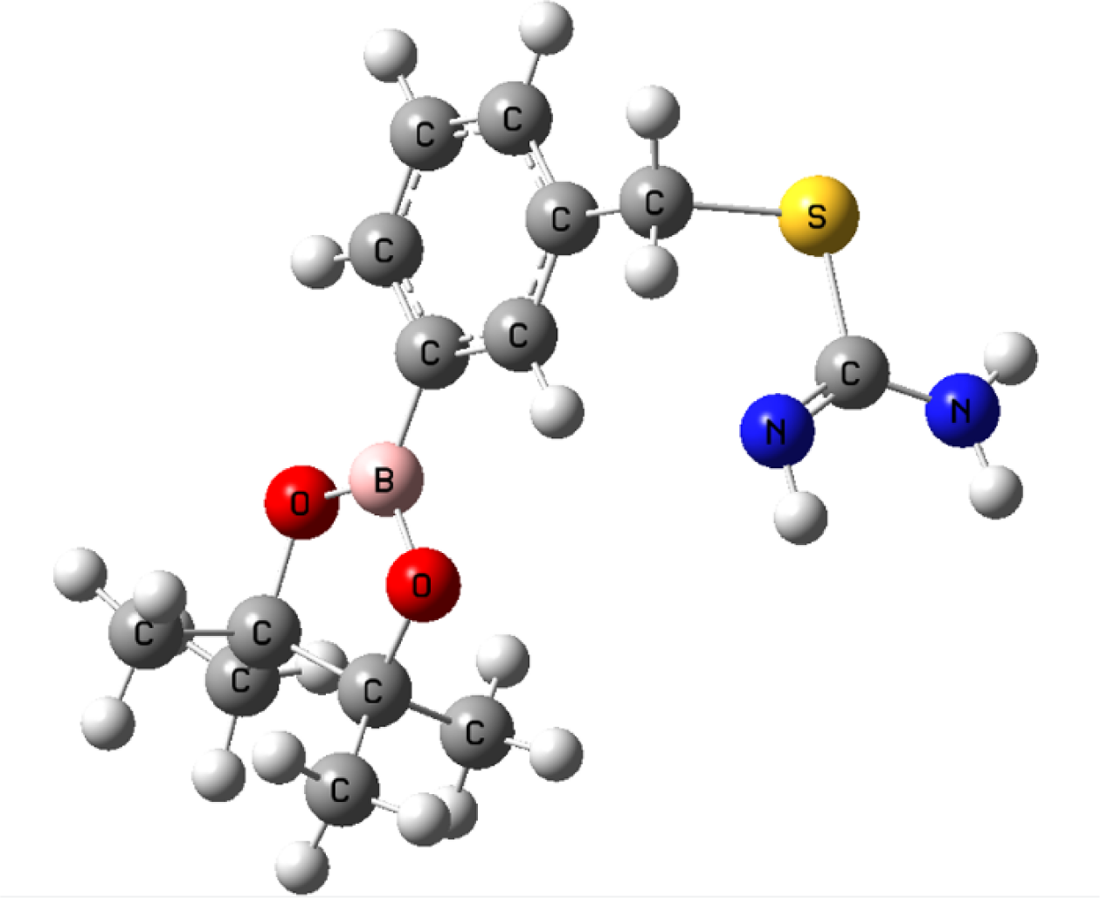
Optimized geometry of ligand *C_14_H_21_BN_2_O_2_S*

#### 5. Frequency calculation for all optimized geometries

To ensure that all the model geometries are fully optimized, frequency calculations were performed on the optimized ligand, protein, and protein-ligand complex using DFT method (B3LYP/LANL2DZ). Result shows that there is no imaginary frequency present in all three model geometries and hence they are the energy minimised structures.

#### 6. Single point energy calculation for all optimized geometries

Single point energies were calculated on all the optimized geometries in both gas phase as well as in solvent phase. As biological systems are surrounded by solvents and fluids, we consider water as the solvent. The ligand and protein molecules acting as solute present within the solvent polarise in response to the solvent polarisation. And conversely the molecules of the solvent rearrange themselves and polarise in response to the charge density of the solute. This polarisation induces redistribution of charges between the solute and the solvent until they reach a state of self-consistency which lowers the energy of the whole system and stabilises it.

#### 7. Ligand Binding Affinity

The ligand binding affinity for ligand C_14_H_21_BN_2_O_2_S with protein uPA was calculated using Equation 1 given below

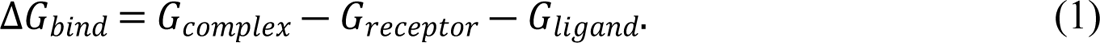

Where

Δ*G*_*bind*_ is the ligand binding affinity

*G*_*complex*_ is the energy of the ligand bound to the binding pocket

*G*_*receptor*_ is the energy calculated for the binding pocket

*G*_*ligand*_ is the energy calculated for optimized ligand

All these G values are calculated in a.m.u (atomic mass unit), therefore, to convert it to kcal/mol, it needs to be multiplied with 627.51. Therefore, the equation 1 can also be written as follows:

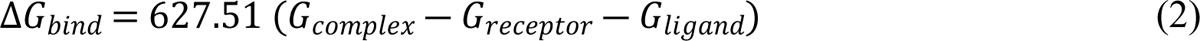

The computed relative energy value (Kcal/mol) for binding affinity of C_14_H_21_BN_2_O_2_S is presented in the Table 1 given below

**Table 1.**
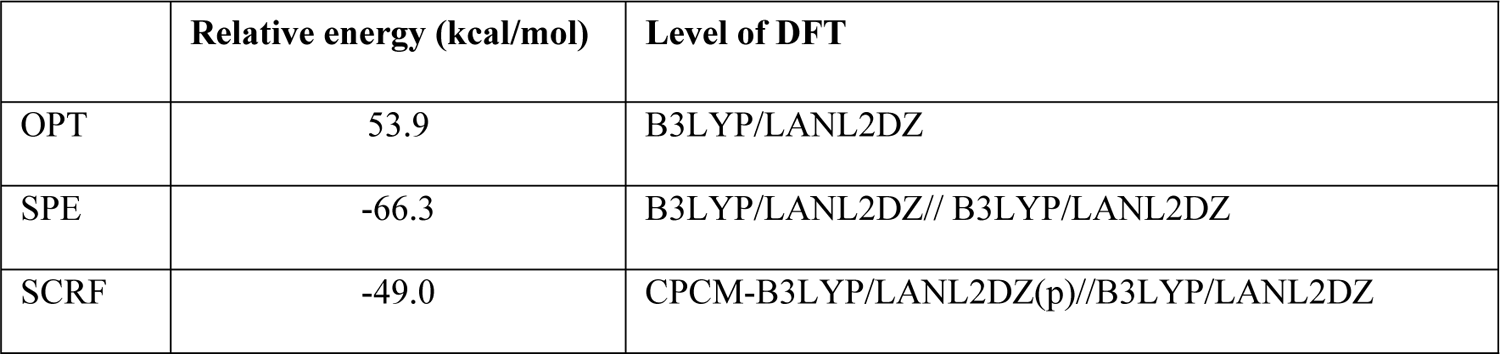
Computed energies in kcal/mol for optimized geometry, Single point energy and self-consistent reaction field for *C_14_H_21_BN_2_O_2_S* bound uPA binding cavity with the methods used as level of DFT

|  | Relative energy (kcal/mol) | Level of DFT |
| --- | --- | --- |
| OPT | 53.9 | B3LYP/LANL2DZ |
| SPE | -66.3 | B3LYP/LANL2DZ// B3LYP/LANL2DZ |
| SCRF | -49.0 | CPCM-B3LYP/LANL2DZ(p)//B3LYP/LANL2DZ |

The results indicate that the binding interaction between C_14_H_21_BN_2_O_2_S and uPA is stable/feasible as the relative energy is exothermic with the release of −49.0 kcal/mol energy.

### D. Pharmacophore modelling

The intention for building a pharmacophore model was to show a concept which explains the importance of different selected pharmacological features in the test compounds and not to optimize the inhibitory effect of these test compounds that act as inhibitors.

All seven are chosen as the template for generating a pharmacophore model based on their high potency and selectivity. The compounds were aligned by a flexible alignment method and used as a template for the random selection of favourable pharmacophoric features. Selection of test set and activity Cut-off 263 boronic acid-derived inhibitors against uPA in humans with known IC_50_ values were selected from the Binding database (BDB). Compounds with unknown IC_50_ values were removed from the test set. Furthermore, the seven test compounds had IC_50_ values lower than 69µM, which makes them fit for activity cut-off. The compounds in the test data with 69µM were considered active compounds, while the compounds above 69µM were considered inactive compounds. Out of the total of 263 compounds, 63 lie in the active compounds, which have IC_50_ values equal to or less than 69µM and 200 compounds lies in the inactive compounds, which have IC_50_ values greater than 69µM, which is the cut-off value as stated before. The model was built through random selection of the different pharmacophoric features and removal of those descriptors that had no effect on the differentiation of active from inactive. The selected pharmacophoric features were revised through modifications or by altering the Gaussian radius so that the maximum number of actives could be selected as hits by the generated pharmacophoric model. The pharmacophore model for the seven ligands was 78% accurate, suggesting that the generated model in this analysis is well-predicted, making it efficient in distinguishing between active and inactive compounds, which indicates it is able to classify actives as hits selectively.

In conclusion, the model shown in figure 15 is finalised for the selected test set of 263 compounds that were able to select all active compounds as hits except for 13 active compounds. The pharmacophore model selected for screening the boronic acid derivative inhibitors of uPA comprises five distinct features that are:

- F1 Aromatic hydrophobic ring (Aro-Hyd),
- F2 Hydrogen bond donor and metal ligator and cation hydrogen bond acceptor [Don&ML&(Cat|Acc)],
- F3 Hydrogen bond donor and metal ligator and cation hydrogen bond acceptor [Don&ML&(Cat|Acc)],
- F4 Metal ligator and hydrogen bond acceptor and cation and hydrogen bond donor [ML&(Acc|Cat|Don)],
- F5 Metal ligator and hydrogen bond acceptor and cation and hydrogen bond donor [ML&(Acc|Cat|Don)]. These selected pharmacophoric features have a radius within the range of 1.0-2.0°A. The sensitivity and specificity were also calculated for the generated pharmacophore model, which signifies the correctness of the model. By putting the values within the present equations, the following solutions were generated:

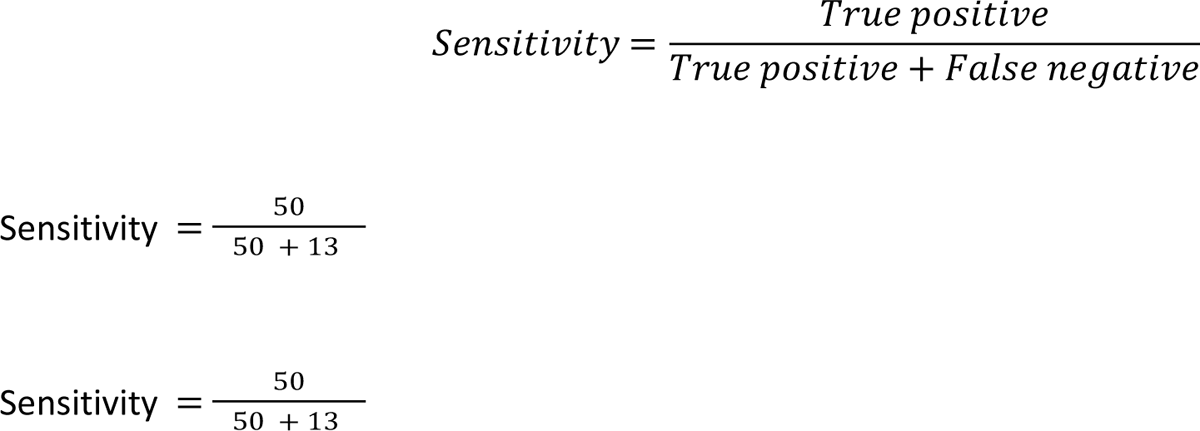

Sensitivity = 79%

The model has 79% sensitivity to distinguishing between active and inactive test compounds.

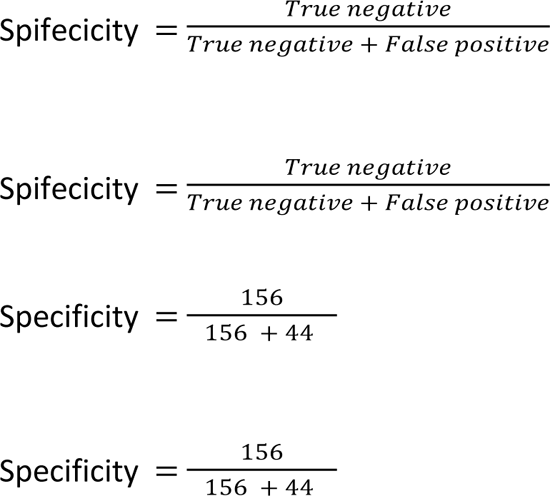

Specificity = 78%

The model has 78% specificity towards active compounds as hits.

**Figure 15.**
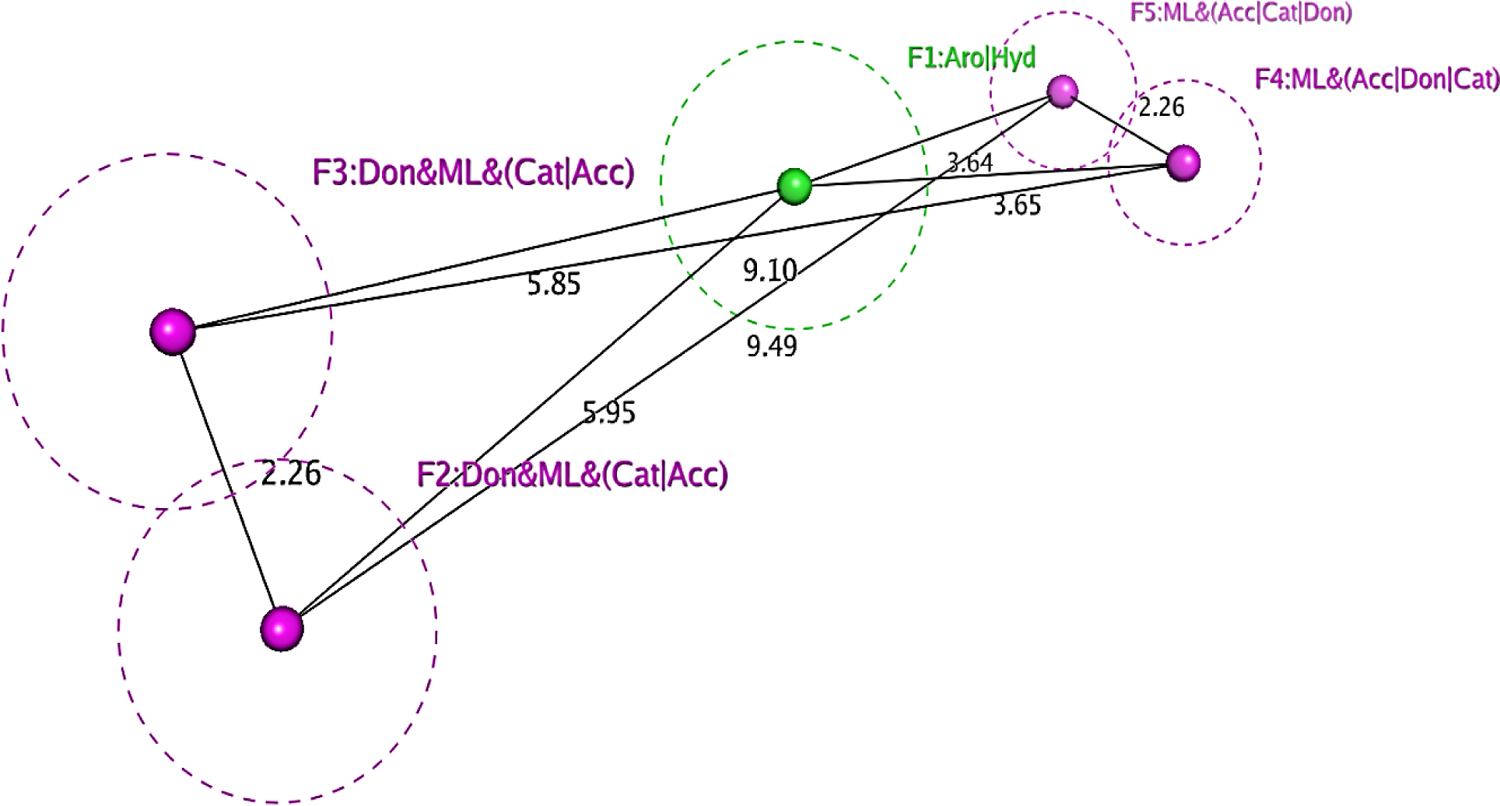
Statistically significant (19% True positive and 59.3% True negative rate) boronic acid derivative inhibitors pharmacophore model obtained using docking conformations of seven inhibitors aligned with flexible alignment used as a template. The pharmacophore consists of four cationic hydrogen bond acceptors and two cationic hydrogen bond donors with one aromatic ring.

## 5. Discussion

Computational quantum mechanical techniques are unable to produce accurate results for binding free energies specifically for ligands and proteins in a solvent environment that mimic the real situation. Consequently, the quantum mechanical techniques would be appropriate for prediction purposes instead of the precise estimation of binding free energies for the ligands and proteins. The development of inhibitors containing moieties which can interact with different uPA sub-sites with high selectivity, potency, and improved pharmacokinetic properties are the main challenges at this stage. It is hoped that the results presented here should stimulate combining experimental and theoretical works for developing uPA inhibitors in cancer treatment through a better understanding of the binding interaction of uPA and its inhibitors. Structure-based technique molecular docking was used to produce a protocol for validation of the previously present binding interactions between seven ligands (C_8_H_11_BN_2_O_2_S, C_8_H_11_BN_2_O_2_S-a, C_14_H_21_BN_2_O_2_S, C₁₈H₃₁N₃O₂S, C_9_H_9_F_3_N_2_S, C₁₁H₁₈BNO₂S and C_9_H_10_N_2_O_2_S) and uPA and ligand-based techniques were combined with them for demonstrating the importance of significant descriptors for optimum biological activity at receptor site by the inhibitor. Molecular docking simulations were performed to hypothesise the binding activity with the help of two software. The results generated were analysed not only based on the calculated score but the residues involved in the binding as well. Ligand C_14_H_21_BN_2_O_2_S was chosen as the most suitable inhibitor among seven, with scores of −3.2481 kcal/mol with MOE and 46.4523 kcal/mol with GOLD. C_14_H_21_BN_2_O_2_S showed interactions with receptor amino acid residues GLY219, SER190 and ASP189 with Sulphur and amino group generated by MOE and GLY219, GLY216, and ASP189 with Sulphur, amine and amino group generated by GOLD. Computational Quantum mechanical studies were applied based upon the electron density of uPA to find the binding energy of active ligand against receptor uPA using hybrid functional B3LYP in combination with LANL2DZ of Density Functional Theory (DFT) as a basis set on the selected model of the active site of uPA. A −2 charge is present on ASP189 of the binding cavity throughout the simulations. From the computational analysis of the calculated values Geometric optimization (opt) = 53.9, Single point energy (SPE) = −66.3 and Self-consistent reaction field (SCRF) = −49.0, it is concluded that uPA shows better binding with ligand when there is a negative two charge on it ASP189 amino acid residue in the binding pocket.

A pharmacophore model was designed because of significant descriptors. These descriptors can be used to search for compounds that may act as efficient inhibitors that fit the model, as these features are characterised as essential regarding the biological activity of inhibitors against uPA. The biological activity was correlated with the effect of 3-dimensional (3D) properties of ligands. It was concluded from the results of QSAR that in uPA, two hydrogen bond donor groups at a distance of 2.2 °A apart were considered favourable for activity. There is an aromatic hydrophobic ring at a distance of 5.95 °A and 5.85 °A from the two hydrogen bond donor groups and at a distance of 3.65 °A and 3.64 °A from the two metal ligator hydrogen bond acceptor groups. Also, the two metal ligator hydrogen bond acceptor groups are situated at 2.2 °A apart from each other. The designed model shows 79% Sensitivity, 78% Specificity and 51% calculated MCC. This model was tested for a test set of 263 boronic acid-derived inhibitors against uPA to predict the accountability of the model by judging how well it can differentiate between active and inactive compounds with a specific activity cut-off. This model can further be tested for liability through experimental methods.

## 6. Materials and Method

### A. Prediction of the protein binding pocket

The protein data bank (PDB) provided the structure chosen for binding site prediction. The PDB contains multiple crystal structures for the same protein, and choosing the best one relies on the type of study. The criteria used to choose the crystal structure uPA(1W10)(16) was

- Resolution of the crystal structure which is claimed to have a resolution of 2.00 A
- A reduced R-factor of 0.190
- The Clash score is only 4, and there are no Ramachandran outliers.

DoGsite scorer and RaptorX are the two non-commercial tools used for the prediction of the binding site for uPA. A PDB file of uPA that was obtained from the Protein Data Bank was used as the input file and docked in various directions. DoGsite scorer predicted nine binding pockets, the second predicted pocket (P 1) with the highest drug score (0.67) was chosen because it included important amino acid residues for the inhibition of uPA (17).

RaptorX needs the protein’s FASTA sequence as an input file for the prediction of the binding pocket. With various multiplicity scores, four predicted binding pockets for the target protein uPA were predicted, and the pocket with the highest multiplicity score was selected. The amino acid residues within the predicted binding pockets were visualised using PyMOL (18). The residues were labelled as H46, D192, S193, C194, Q195, G196, S198, V216, S217, W218, G219, G221, C222 (19).

### B. Molecular docking

Two software tools, molecular operating environment (MOE) and genetic optimization for ligand (GOLD) suit, were used for docking studies. The DOCK module of MOE was used to generate ten poses for each of the seven test ligands, and the steps for the ligand conformational search were placement, scoring, and refinement by energy minimization under a defined force field. The scoring function used was LondonDG (20) in combination with the placement method Alpha triangle for docking optimization protocol, and the forcefield applied for energy minimization was AMBER99 (21). Genetic optimization and ligand binding (GOLD) suit version 5.6.1 was used to generate ten poses for each of the seven test ligands. The genetic algorithm that implements quantitative prediction of binding energies was used to dock the ligand within the target protein uPA with partial flexibility (22). Ligands were docked using the scoring function GoldScore into the binding pocket of the target protein with PDB ID: 1W10 with resolution 2°A. X, Y and Z coordinates were used for the allocation of the binding site in the docking studies within a 15°A radius (23). The generated poses were analysed through MOE to select the most favourable conformation based on the binding interaction between the ligands and the residues of the binding pocket.

### C. Quantum mechanical studies/calculations

A molecular visualisation tool, Swiss-Pdb viewer, was used to extract the binding pocket of uPA from within the whole biomolecule, which was utilized for the generation of model complex geometry (24). The Quantum mechanical (QM) calculations were performed by Density functional theory (DFT) (25) method to find the binding affinity of the test ligands within the binding pocket of the target receptor protein uPA using the commercial software GAUSSIAN 09 (26). Binding energies, single point energies, self-consistent reaction field (SCRF) and frequencies for the molecular complex were calculated, and the results generated were observed with GaussView (27)(28). The extracted model of the binding pocket with bound ligand C_14_H_21_BN_2_O_2_S was passed through a series of clippings and modifications by trimming the amino acid residues ASP189, HIS99, SER214 and TRP215 at their alpha carbon (α − carbon) atom positions, and hydrogen atoms were added to satisfy the valency. The first step was hydrogen optimization, followed by geometry optimization of the model complex with hybrid density functional method B3LYP in combination with a basis set LANL2DZ. For both geometries, some of the atoms at the alpha carbons of amino acid residues ASP189, HIS99, SER214 and TRP215 are fixed. For ligand model geometry optimization, ligand C_14_H_21_BN_2_O_2_S was extracted from the X-ray crystal structure and optimised using the B3LYP/LANL2DZ level of DFT studies. The protein binding pocket geometry model was obtained by eliminating the ligand structure and optimised with the same level of DFT, B3LYP/LANL2Z. The optimised geometries were used in the calculation of the single point energies in both gas and solvent phases using the B3LYP/LANL2DZ level of DFT. To ensure that all the model geometries are fully optimized, frequency calculations were performed on the optimized ligand, protein, and protein-ligand complex using the DFT method (B3LYP/LANL2DZ). Stuttgart Dresden’s (SDD) effective core potential basis set was used for the calculation of the single point energy in both the gas phase (vacuum) and the solvent phase (water) for all the optimized geometries. The electronic energy of the model complex geometry is calculated as the single point energies. Using Molden, GAUSSIAN output files are visualised, showing the atomic density, electron density, molecular density, and molecular orbitals (29).

### D. Pharmacophore model

For the target protein, uPA pharmacophore queries were generated by using MOE software. The model was built on the basis of those descriptors that could efficiently differentiate the active from the inactive. The selected pharmacophoric features were revised through modifications or by altering the Gaussian radius so that the maximum number of actives could be selected as hits by the generated pharmacophoric model (30)(31).

## 7. Acknowledgments

All the praises be to Almighty Allah the most compassionate and the most merciful, who has bestowed upon me the power and ability to think and grow, empowering me to play my role in conveying a little share of my knowledge. I will forever be grateful to my dear father and loving mother who helped me get through my research phase and for their love and emotional support, for always encouraging me to keep going no matter what. I would like to menion my husband for supporting me through the end of my research phase and always being there for me. Moreover, I would like to add, I would not have accomplished what I have without the constant support of my generous supervisor Dr. Mehak Rafiq on whom I relied for guidance and help at any time. I am much obliged to my guidance committee members Dr. Rehan Paracha and Dr. Uzma Habib who have always been available for their humble assistance at various stages of my study and provided me with their valuable feedback and opinion.

